# *Fgf10* mutant newts can regenerate normal limbs despite severe developmental hindlimb defects

**DOI:** 10.1101/2023.05.23.541496

**Authors:** Miyuki Suzuki, Akinori Okumura, Yuki Shibata, Tetsuya Endo, Machiko Teramoto, Akane Chihara, Kiyokazu Agata, Marianne E. Bronner, Ken-ichi T. Suzuki

## Abstract

In the amniote limb, FGF10 is essential for limb bud initiation and outgrowth. However, whether this function is broadly conserved in tetrapods and/or involved in adult limb regeneration remains unknown. To tackle this question, we established an *Fgf10* null mutant line in the newt *Pleurodeles waltl* which have amazing regenerative ability. While *Fgf10* mutant forelimbs develop normally, the hindlimbs exhibit severe digit reduction, fail to ossify the zeugopod, and downregulate FGF target genes like *Sall1, Runx1* and *Hoxa11/d11*. Despite these developmental defects, *Fgf10* mutants were able to regenerate near-normal hindlimbs. Together, our results suggest an important role for *Fgf10* in hindlimb digit formation and zeugopod ossification during development, but little or no function in regeneration, suggesting that different mechanisms operate during limb regeneration versus development.

## Results and Discussion

*Fgf10* is essential for initiation and outgrowth of the limb bud of amniotes ^1,2^. Activated by *Tbx5/4* from lateral plate mesoderm ^3-5^, *Fgf10* is expressed throughout the limb mesenchyme, where it stimulates *Fgf8* expression in the apical ectodermal ridge (AER). Genetic ablation of *Fgf10* in mice is lethal and leads to severe truncation of both fore- and hind-limbs as well as developmental defects of epithelial components in multiple organs including the lung. Whether this function is evolutionary conserved throughout tetrapods remains unknown.

Many urodeles exhibit different patterns of FGF gene expression from those observed in amniotes. In axolotl, for example, *Fgf8* is expressed in the mesenchyme rather than in epidermis ^6,7^ suggesting that *Fgfs* may work differently in amniote than urodele amphibian limb development. Moreover, in contrast to amniotes, many urodeles have the ability to regenerate their limbs throughout adulthood. While it has been suggested that adult limb regeneration may recapitulate development, it is unclear whether or not FGFs function in similar ways during development and regeneration.

To address the role of FGF10 in limb development and regeneration, we have tested its function in *Pleurodeles waltl* (Iberian ribbed newt) due to its remarkable ability to regenerate organs throughout adulthood. This coupled with the efficiency of CRISPR-Cas9 mediated Knock-out (KO) and availability of transcriptomic and genomic information ^8-11^ make this newt an ideal model for exploring similarities and/or differences between limb development and regeneration. As a first step to examine the function of FGFs in limb development, we detected the expression patterns of *Fgf8* and *Fgf10* transcripts using hybridization chain reaction (HCR) in limb buds. Similar to the result observed in the axolotl, both *Fgf8* and *Fgf10* were expressed in the limb mesenchyme in *Pleurodeles waltl* during development (Fig. 1A)^12,13^. *Fgf8* was also expressed in mesenchyme of the limb blastema (fig. S1). Bulk-RNA seq confirmed that *Fgf8* and *Fgf10* were expressed in both fore- and hind-limbs (Fig. 1B).

**Fig. 1.**
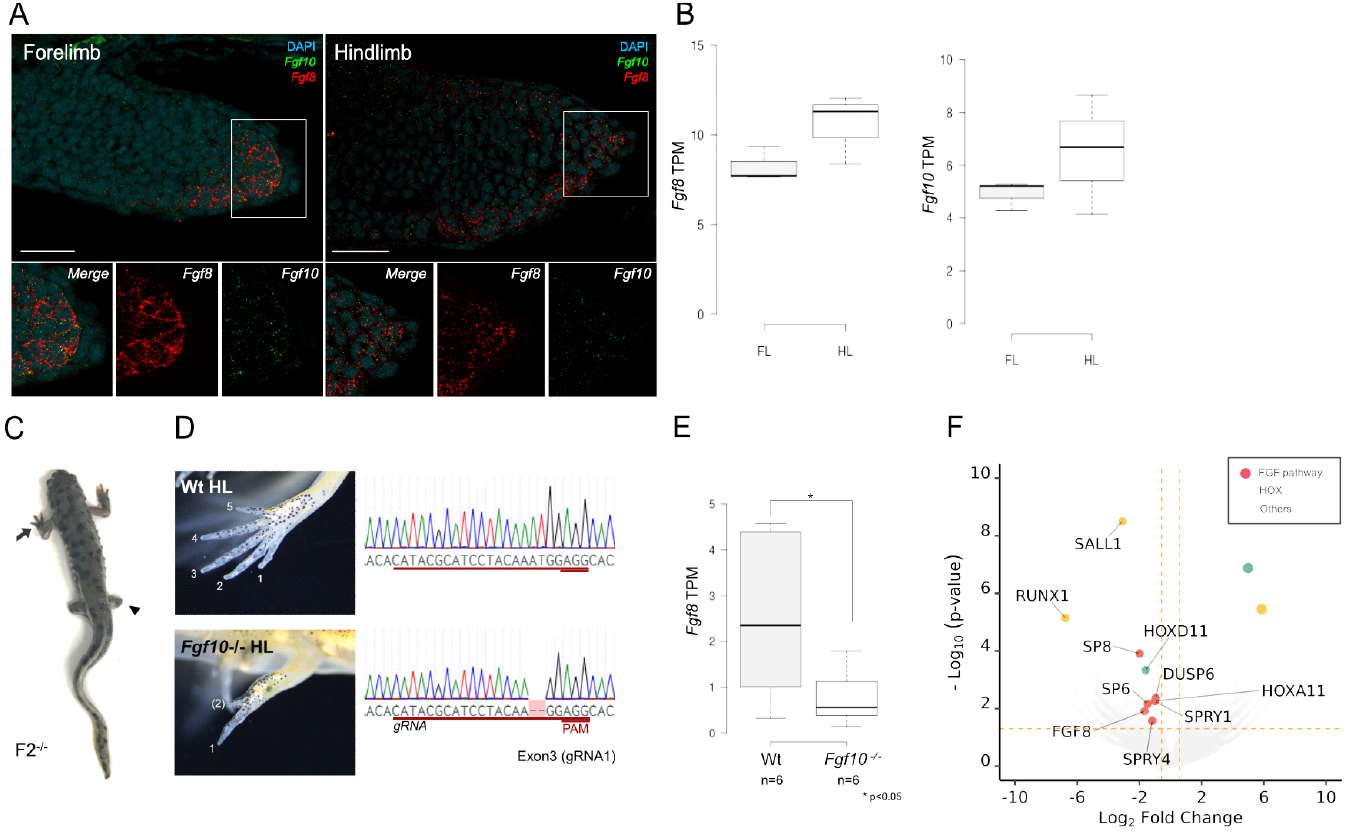
*Fgf10* mutant has defect only in hindlimb. (**A**) *in situ* HCR image of *Fgf8* and *Fgf10* in fore- and hindlimb bud. Scale bar = 50 µm. (**B**) *Fgf8* and *Fgf10* mRNA expression level in forelimb (FL) and hindlimb (HL) (n=3 biological replicates) (**C**) The phenotype and genotype of *Fgf10* null mutant (F2). Arrow and arrowhead show normally developed forelimb and defective hindlimbs, respectively. (D) Phenotype and sanger sequence chromatograms of wild type and *Fgf10* F2 mutant. The sequence of gRNA, PAM, and deletion are indicated by red underlines and dash. (**E**) *Fgf8* expression in *Fgf10* mutant hindlimb bud. * p<0.05 (**F**) Differential gene expression analysis of *Fgf10* mutant hindlimb bud (n=3 Wt; 6 *Fgf10* mutant; *p < 0.05).

To test the functional role of *Fgf10*, we designed gRNAs to *Fgf10* and injected Cas9 ribonucleocomplex into a one-cell-stage embryo. In contrast to results in mice, the forelimb of *Fgf10* crispants developed normally; however the hindlimb was defective with a digit number reduced to one or two (fig. S2). This phenotype was reproduced with multiple gRNAs (fig. S3), and the mutation rate was more than 93% in both fore- and hind-limb (fig. S4) suggesting that the observed difference in phenotype between the two limbs was not due to mosaicism. Furthermore, we generated an *Fgf10* F2 null mutant by crossing F1 heterozygotes with a 2bp deletion in the target site. These exhibited the same defect that was restricted to the hindlimb while forelimb was normal (Fig. 1CD). One possible explanation for the differences between forelimb and hindlimb is that other *Fgfs* may function redundantly with *Fgf10* in the forelimb.

To investigate the expression of downstream genes, we collected six hindlimb buds individually and performed RNA-seq analysis. As expected, *Fgf8* was significantly downregulated in the *Fgf10* mutant (Fig. 1E). Other *Fgf* pathway genes, *Sp6, Sp8, Spry1*, and *Spry4* were also downregulated (Fig. 1F). The most significantly downregulated gene was *Sall1. Sall4* is considered to be a putative target of thalidomide which leads to limb defects in humans ^14^ and *Sall1* has been suggested to be functionally redundant *Sall4* in several developing organs ^15^, raising the intriguing possibility that this may contribute to defects in limb development. In addition, *Hoxa/d11* and *Runx1* were significantly decreased (Fig. 1F).

*Hoxa11* and *Hoxd11* are expressed in the zeugopod region of the limb and their deletion results in zeugopod malformations ^16^. The transcription factor *Runx1* is essential for osteogenesis ^17^. From micro CT analysis of adult *Fgf10* heterozygotes and null mutants (1-year post-fertilization), we found that heterozygotes had proper osteogenesis in the stylopod, zeugopod, and autopod (Fig. 2Aa) while the null mutant had fewer digits and lacked fibula or tibia ossification in the zeugopod region (Fig. 2Ab). Alizarin red/Alcian blue staining of *Fgf10* crispant juveniles also showed that even while the digits and stylopod completed ossification, zeugopods were slow to ossify and again the fibula or tibia were not ossified (Fig. 2B). Based on these phenotypes and RNA-seq analysis, we suggest that *Fgf10* plays two roles: first in autopod patterning by coordinating *Fgf* pathway and *Sall* genes; second in zeugopod ossification through *Hox11* genes and *Runx*.

**Fig. 2.**
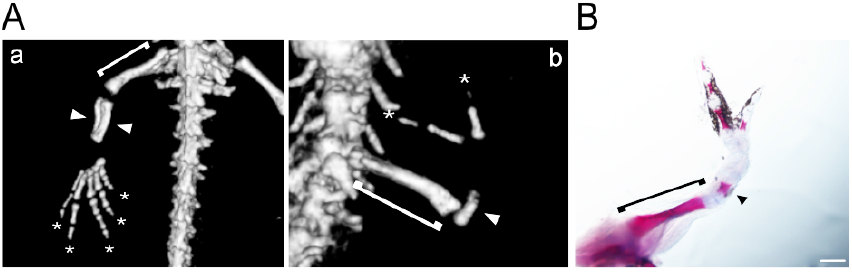
*Fgf10* mutation causes zeugopod malformation in hindlimb. (**A**) microCT image of the hindlimb of adult *Fgf10* mutant. (a) Control. *Fgf10* hetero mutant which shows normal hindlimb development. (b) *Fgf10* null mutant. The bracket, arrowhead, and asterisk show stylopod, fibula/tibia, and digits, respectively. (**B**) Bone staining of *Fgf10* crispant. The arrowhead shows no ossification of fibula or tibia in zeugopod. Scale bar = 1mm.

Given that limb development and limb regeneration share many common properties, we next examined the role of *Fgf10* in limb regeneration. To this end, we amputated either the forelimb or hindlimb of *Fgf10* crispant animals and the hindlimb of F2 null mutants (1 or 2.5 month post fertilization, respectively). The forelimb of *Fgf10* crispants develops normally (4 digits). After amputation, the forelimbs regenerated normally to the original number of digits (fig. S5). In contrast to the developing forelimb, the developing hindlimb exhibited profound developmental defects, forming only 1 or 2 digits. Surprisingly, after amputation, rather than recapitulating the original morphology, these limbs regenerated additional digits in *Fgf10* F2 null mutants (Fig. 3AB). Whereas the *Fgf10* F2 null mutants have 0 to 2 digits during hindlimb development, amputation of their limbs increased the number of digits in more than half of the mutants, and 28% of mutant completed regenerated identical to the wild type with 5 digits (n=5; Fig. 3AB).

**Fig. 3.**
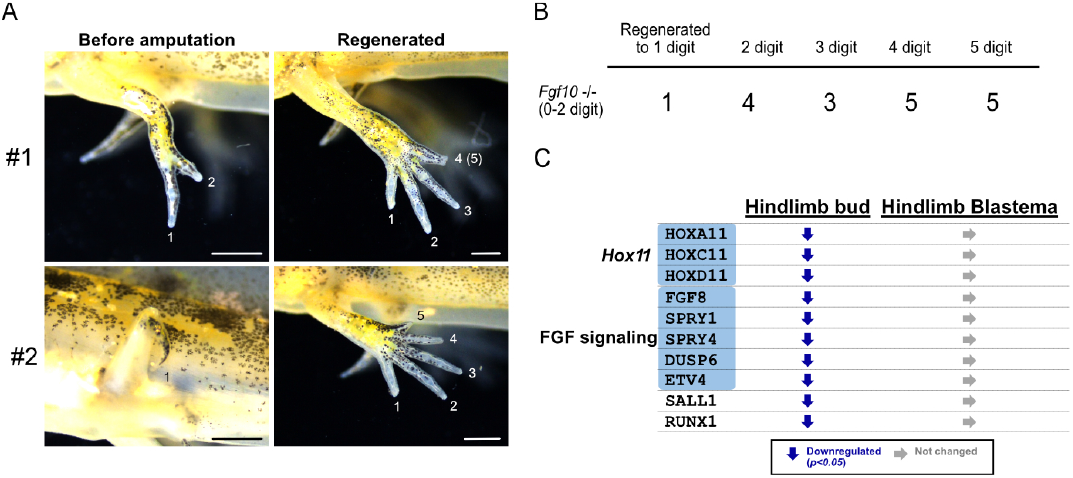
Defected hindlimb in *Fgf10* mutant was recovered by regeneration. (**A**) Hindlimb regeneration of *Fgf10* mutant from two different individuals. Scale bar = 1 mm. (**B**) Summary of restored hindlimb in *Fgf10* null mutant (n=18). (**C**) Summary of differential expression analysis of limb development-related genes. Down and right arrow indicates significantly downregulated (p<0.05) and no difference in Fgf10 null mutant limb bud or blastema compared with wild type.

To examine downstream gene expression changes, hindlimb blastemas from wild-type (n=3) or *Fgf10* null mutants (n=6) were individually sampled and processed by RNA-seq. The expression levels of *Fgf10* regulated genes were found to be comparable between the wild type and mutant blastemas, suggesting that downstream gene expression was restored (Fig. 3C, fig. S6). Other limb development-related genes including the *Fgf* signaling pathway were also similar between wild type and mutant blastemas. These results suggest that hindlimb regeneration does not recapitulate development but rather occurs by a different mechanism.

Finally, we examined ossification of the zeugopod during mutant limb regeneration. Micro CT imaging of the regenerated limb of *Fgf10* null mutant adults revealed normally ossified fibula and tibia (Fig. 4). Thus, the limbs recovered not only autopod patterning but also ossification of the zeugopod, suggesting that the regenerated *Fgf10* mutant hindlimb is comparable to the wild type. These results suggest that *Fgf10* is not essential for forelimb or forelimb development and hindlimb regeneration, but is required for hindlimb development.

**Fig. 4.**
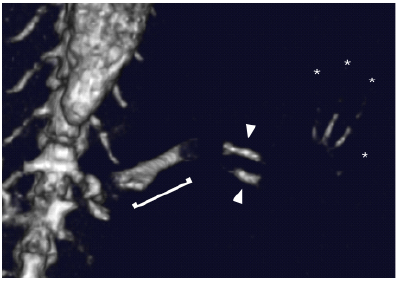
Zeugopod malformation in hindlimb was recovered by regeneration. microCT image of regenerated hindlimb of *Fgf10* null mutant. The bracket, arrowhead and asterisk show stylopod, fibula/tibia, and digits, respectively.

Taken together, the results suggest that developing and regenerating urodele amphibian limbs differentially utilize *Fgfs*. While *Fgf10* is required for hindlimb development, it is not required for regeneration or for either development or regeneration of the forelimb. Our results in the hindlimb suggest that the *Fgf* pathway contributes to autopod patterning via *Sall* and to zeugopod ossification through *Hox11* and *Runx* genes. However, for hindlimb regeneration, there was no loss of *Fgf8* and other *Fgf* pathway members in *Fgf10* mutants. Notably, this demonstrates that development and regeneration utilize different *Fgf8* induction mechanisms (Fig. 5). Reciprocal and feedback loop regulation between *Fgf10* expressed in mesenchyme and *Fgf8* expressed in AER is essential for vertebrate limb development^1,2^. However, both of these *Fgfs* are expressed in the mesenchyme of urodeles^6,18^. One important difference between development and regeneration is the presence of nerves in the blastema, particularly since nerve-derived factors play a critical role in regeneration. In the axolotl, grafting of *Fgf8* with other growth factor beads into wounded skin ^19^ or over-expression of *Fgf8* plus rerouted nerves ^18^ can induce an ectopic blastema and limb outgrowth in the upper arm. These findings together with the present results suggest the possibility that direct induction of *Fgf8* in the mesenchyme by nerve derived factor(s) rather than *Fgf10* is the key to limb regeneration in urodeles (Fig. 5).

**Fig. 5.**
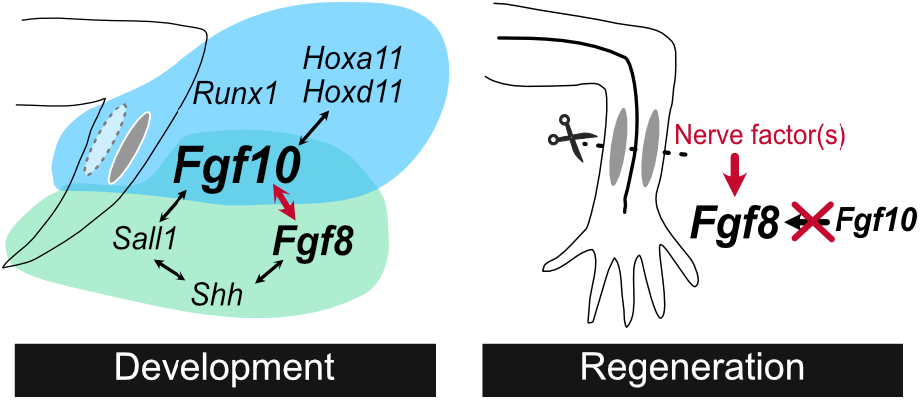
A model for FGF regulation in urodele hindlimb development and regeneration.

Our findings highlight both similarities and differences between limb development and regeneration. Bryant and colleagues have proposed that the early phase of limb regeneration may involve steps that are regeneration-specific. Subsequently, there is a gradual transition to an autonomous limb patterning program similar to that utilized during development ^20^. We find that most of the downstream genes required for limb development, including *Fgf8*, do not differ between *Fgf10* null and wild type blastema. Accordingly, we suggest that it is possible that only a limited number of genes, including *Fgf10*, have differential effects at early stages of regeneration whereas later events in regeneration parallel those in play during development. Importantly, despite the fact that *Fgf10* mutant hindlimbs display marked abnormalities, our results show that the hindlimb has no “developmental memory” and is able to regenerate a normal limb from a developmentally defective one.

## Materials and Methods

### Animals

The Iberian ribbed newts (*Pleurodeles waltl*) used in this study were obtained from a breeding colony at Tottori University and Hiroshima University. For anesthesia before limb regeneration and samplings, MS-222 (Sigma, St. Louis, MO, USA) was used at a final concentration of 0.02%. The developmental stages were defined according to previous methods ^21^. The forelimb was amputated in the middle of the zeugopod at 1-month post-fertilization (mpf). The hindlimb was amputated at 2.5 mpf in the middle of the zeugopod or stylopod if its zeugopod is severely truncated. Blastemas were collected at 10 days post-amputation (dpa) and allowed to regenerate for 1 month before observation. Animal rearing and treatments were performed and approved in accordance with the Guidelines for the Use and Care of Experimental Animals and the Institutional Animal Care and Use Committee of the National Institute for Basic Biology and California Institute for Technology.

### *In situ* hybridization chain reaction (HCR)

Embryos (st. 34-35 for forelimb, st. 39-41 for hindlimb) and blastema tissue were collected under the anesthesia and embedded in low melt agarose and/or Tissue-Tek OCT compound (SAKURA, Nagano, Japan) and immediately frozen in liquid nitrogen. Tissues were sectioned at 14μm. HCR probes (*Fgf8* and *Fgf10*) were purchase from Molecular Technologies and Molecular Instruments (Los Angeles, CA, USA) and *in situ* HCR was performed following the manufacturer’s instructions.

### sgRNA design and Cas9 injection

*PwFgf10* sequence was obtained from a previous report (M0204228_PLEWA04^10^) and compared with other species to define exon-intron boundary (fig. S3B, C). Three sgRNAs (gRNA1-3) were designed at exon 3 in the same location as used for the *Fgf10* crispant mouse ^22,23^, and one sgRNA was designed at exon 1 downstream of the ATG site. *In vitro* transcription of sgRNA and Cas9 ribonucleocomplex injection was performed as previously described ^9^.

### RNA sequencing and differential gene expression analysis

The fore- and hind-limb buds were collected at st.33-34 and st.39-40 and pooled separately. For differential gene expression analysis in limb development, the hindlimb bud of wild type or *Fgf10* F2 null mutants were collected individually (n=6, each) at st. 39-40. For limb regeneration, fully grown blastemas (10 dpa) were collected individually (n=3 for wild type, n=6 for *Fgf10* mutant). Each sample was soaked in RNAlater (Invitrogen, Waltham, MA, USA) at 4°overnight, and stored at -80° until extraction. Total RNA was extracted and purified using NucleoSpin (Takara Bio, Shiga, Japan), following the manufacturer’s instructions. The RNA-Seq of normal fore- and hind-limb bud was conducted by Macrogen Japan corp (Kyoto, Japan) and the RNA-Seq libraries of blastemas in wild type and *Fgf10* null mutant were generated using the Illumina stranded mRNA Prep kit (Sandiego, Cam USA) and sequenced in GENEWIZ (South Plainfield, NJ, USA). Paired-end sequencing of each 150 bp end of cDNA was performed with the Hiseq2000 sequencing system (Illumina, San Diego, CA, USA). The reads were mapped to the reference transcript database (PLEWA04_ORF.cds.fa^10^) using Salmon ^24^ and differential expression analysis was performed by edgeR GLM ^25^. The volcano plot and box plot were generated by ggVolcanoR ^26^ and BoxPlotR ^27^. The data have been deposited with links to BioProject accession number PRJDB15083 in the NCBI BioProject database (https://www.ncbi.nlm.nih.gov/bioproject/).

### Genotyping

For *Fgf10* crispant mutation analysis, both fore- and hind-limbs were collected individually for amplicon sequencing analysis as previously described ^28^. For the *Fgf10* F2 null mutant, genomic DNA was extracted from the tail tip or limb using DNeasy Blood and Tissue Kit (Qiagen, Hilden, Germany), and sequenced by the Sanger sequencing system.

### Micro CT and bone staining

MicroCT image of *Fgf10* F2 mutant (1-year-old adult) was obtained using Rigaku R_mCT2 (Rigaku, Tokyo, Japan). For bone staining, juveniles of *Fgf10* crispant (2-4 mpf) were fixed in 10% formalin and stained with Alizarin red/Alcian blue following a previously reported protocol ^29^. The skin was removed before observation.

## Acknowledgments

We thank Profs. Toshinori Hayashi in Hiroshima University and Takashi Takeuchi in Tottori University for providing *P. waltl* newts. We thank the Beckman Institute Biological Imaging Facility of Caltech for technical assistance with microscopy experiments.

## Author contributions

Conceptualization: MS, MB, TE, KTS

Methodology: MS, MT, KTS

Investigation: MS, AC, AO, YS, MT, KTS

Visualization: MS, AO, MT, KTS

Funding acquisition: KTS

Project administration: MS, KTS

Supervision: MB, KA, KTS

Writing – original draft: MS, MB, KTS

Writing – review & editing: MS, AC, AO, YS, TE, MT, KA, MB, KTS

## Data and materials availability

All data are available in the main text or the supplementary materials.

## Supplementary Materials

Figs. S1 to S6

