## Supplementaly Fig. S1 to S6 for "*Fgf10* mutant newts can regenerate normal limbs despite severe developmental hindlimb defects"

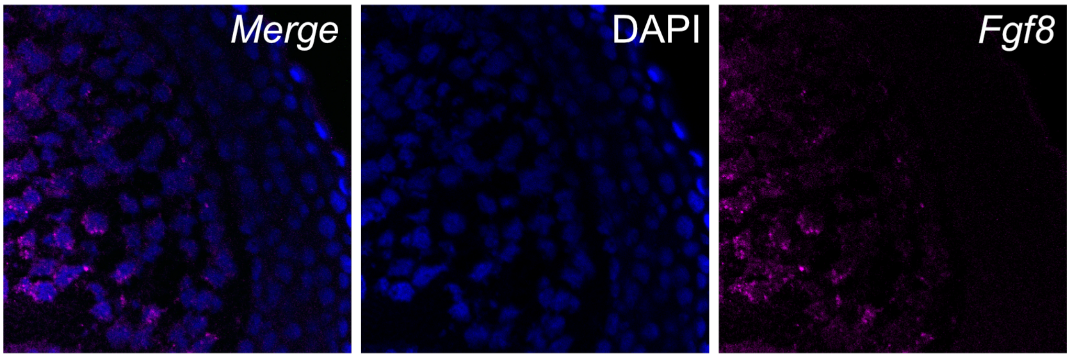


**Supplementary Fig. S1.** *in situ* HCR image of *Fgf8* in forelimb blastema

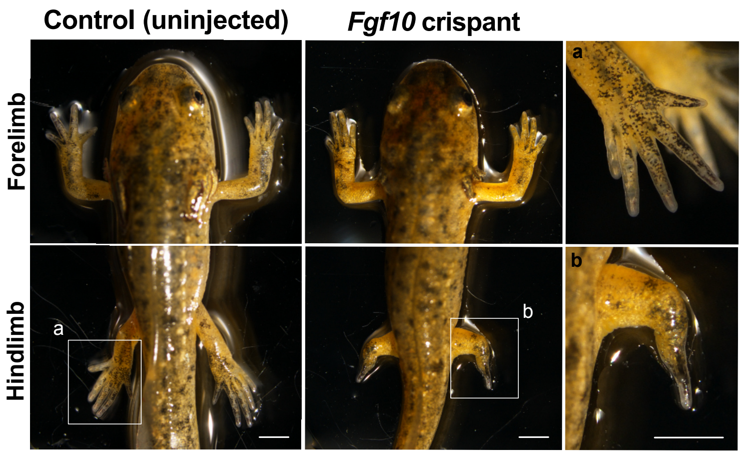


**Supplementary Fig. S2.** The phenotype of *Fgf10* crispant. Scale bar = 2 mm.

A

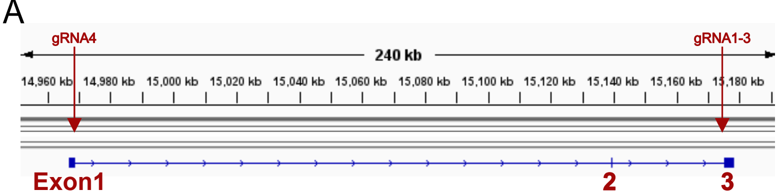


B

```
>Pwfgf10_M0204228_PLEWA04

ATGTGGGGATGGGTACTGAGGAAGGGTGCCTCCGCCTTGTCCCAGCTGCCCTGCCTGGCA
Exon1                          gRNA4
CTGCTGCTCTGGGTGTCGTGCCTGTCGGTCACCTGCCATGACCTGGGCAGGGGCTTGCTG

TCGCCGAGGGTCGCCAATGGCTCGGTGCCTGTGGTGGGCAGACACGCAAGGAGCTACCCG

CACCTCGAGGGAGATGTGCGCCTGAGGAGACTGCTGTGCGTCACCAACTACTTCTTGAAG

ATCGACGCGGACGGCAAGGTCGGAGGGACCACCAAGGAGGACTGCCCGTACAGTGTGTTG
                                                       Exon2
GAGATAACATCCGTGGATGTTGGAATAGTTGCAGTGAGAGGTGTAGAAAGCAAATATTTT

TTAGCAATGAATGAGAAGGGAAAGGTCTACGGCTCGAGAGAATTTACTACAGACTGCGAG
                                    Exon3
CTGAAGGAACGGATGGAGGAAAACAAATACAACACATACGCATCCTACAAATGGAGGCAC
                                                      gRNA1
AAGCAAAGGCAAATGTTTGTGGCCTTAAATGGGAAGGGAAAACCCAAGCGAGGACAGAAA
                                    gRNA2           gRNA3
ACAAAACGGAAAAACACCTCCGCTCACTTTCTTCCGATGCAGATCGACAGACGCTTATAC

TAG
```

C

```
Human           MWKWILTHCASAFPHLPGCCCCCFLLLFLVSSVPVTCQALGQDMVSPEATNSSSSS--FS
Mouse           MWKWILTHCASAFPHLP-GCCCCFLLLFLVSSFPVTCQALGQDMVSQEATNCSSSSSSFS
Axolotl         MCEWVLRKGAAALSQLP-----CLALLLWVSCLSVTCHDLARDMLSPEVAN--------S
Newt            MWGWVLRKGASALSQLP-----CLALLLWVSCLSVTCHDLGRGLLSPRVAN--------G
Zebrafish       MCKWKVTKGASAWFRLS-CLSLPLLLLFLCSALPVACHDTHRAIRAPRGTNS-------S
                *  * : : *:*  :*.       : **:  *...*:*:   : : : . :*       .

Human           SPSSAGRHVRSYNHLQGDVRWRKLFSFTKYFLKIEKNGKVSGTKKENCPYSILEITSVEI
Mouse           SPSSAGRHVRSYNHLQGDVRWRRLFSFTKYFLTIEKNGKVSGTKNEDCPYSVLEITSVEI
Axolotl         SVPVVGRQVRSYKHLEGDVRLRRLLCVTNYFLKIDADGKVSGTTKVDCPYSVMEITSVDV
Newt            SVPVVGRHARSYPHLEGDVRLRRLLCVTNYFLKIDADGKVGGTTKEDCPYSVLEITSVDV
Zebrafish       SSAVVGRHVRSYNHLTGDVRRRKLFSYQKFFLRIDKNGKVNGTKSKDDPYSTLEIKSVDV
                *  .  .**:.*** ** **** *:*:.   ::** *: :***.**.. : *** :**.**::

Human           GVVAVKAINSNYYLAMNKKGKLYGSKEFNNDCKLKERIEENGYNTYASFNWQHNGRQMYV
Mouse           GVVAVKAINSNYYLAMNKKGKLYGSKEFNNDCKLKERIEENGYNTYASFNWQHNGRQMYV
Axolotl         GIVAVKGVYSNYYLAMNEKGRVYGSREFTTDCKLKERMEENKYNTYASYKWRHKQRQMFV
Newt            GIVAVRGVESKYFLAMNEKGKVYGSREFTTDCELKERMEENKYNTYASYKWRHKQRQMFV
Zebrafish       GIVAIKGIQSNYYLAINKKGVVYGARDFGIDCKLIERIEENRYNTYASAEWMNKKKHMFV
                *:**::.: *:*:**:*:** :**:::*   **:* **:*** ****** :* :: ::*:*

Human           ALNGKGAPRRGQKTRRKNTSAHFLPMVVHS----
Mouse           ALNGKGAPRRGQKTRRKNTSAHFLPMTIQT----
Axolotl         ALNGKGTPKRGQKTKRKNTSAHFLPMQIDRHYIR
Newt            ALNGKGKPKRGQKTKRKNTSAHFLPMQIDRRLY-
Zebrafish       GLSANGRPMRAKKTRRKNTATHFLPIPIPIV-----
                .*..:* * *.:**:****::****: :
```

D

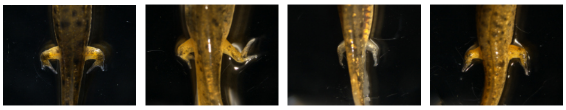

| | Control | *Fgf10* sgRNA_4 | *Fgf10* sgRNA_1 | *Fgf10* sgRNA_2 | *Fgf10* sgRNA_3 |
|---|---|---|---|---|---|
| Normal | 32 | 7 | 11 | 17 | 2 |
| Phenotype | 0 | 6 | 16 | 2 | 17 |

**Supplementary Fig. S3.** Target site and phenocopy of *Fgf10* crispant (A) The IGV image of *Fgf10* genomic region in *Pleurodeles waltl*. *Fgf10* gRNA 1-3 and gRNA 4 were designed at exon3 and exon1, respectively. (B) *PwFgf10* cDNA sequence. Red character and colored underline indicate a start codon and exon 1 to 3. Red highlight and black underline show protospacer adjacent motif (PAM) sequence and gRNA target sequence. (C) Amino acid sequence alignment of the representative sequence of FGF10 compared with four different species (Mouse; AAH48229.1, Human; O15520, Axolotl; ANB78796.1, Zebrafish; AAN62915.1). (D) Confirmation of phenocopy of *Fgf10* crispant using four different gRNAs. The number of individuals who developed normal digit numbers (Normal) and one to two digits in the hindlimb (Phenotype).

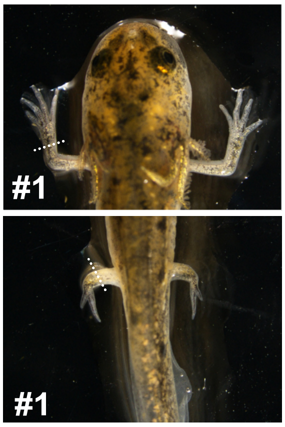


**#1**

| | | Forelimb | Hindlimb |
|---|---|---|---|
| TGGGAAGGGAAAACCCAAGCGAGGACAGAAAACA | Wt | 6.6% | 4.5% |
| TGGGAAGGGAAAAC-------------------A | Δ19 | 63.4% | 10.2% |
| TGGGA-----------------GGACAGAAAACA | Δ17 | 15.2% | 41.5% |
| T-------------CCAAGCGAGGACAGAAAACA | Δ13 | 14.8% | 43.8% |
| | Total reads: | 19,984 | 18,593 |

**#2**

| | | Forelimb | Hindlimb |
|---|---|---|---|
| TGGGAAGGGAAAACCCAAGCGAGGACAGAAAACA | Wt | 4.6% | 5.2% |
| TGGGAAGG----------------ACAGAAAACA | Δ16 | 55.3% | 2.33% |
| TGGGAAGGGA------------GGACAGAAAACA | Δ12 | 40.1% | 92.5% |
| | Total reads: | 15,998 | 15,210 |

**#3**

| | | Forelimb | Hindlimb |
|---|---|---|---|
| TGGGAAGGGAAAACCCAAGCGAGGACAGAAAACA | Wt | 0% | 0% |
| TGGGAAGGGAAA--------------------CA | Δ12 | 100% | 100% |
| | Total reads: | 16,329 | 18,841 |

**Supplementary Fig. S4.** The genotype of fore- and hindlimb of *Fgf10* crispant analyzed by amplicon sequencing. Representative mutant alleles, their occupancy rates, and total read counts are shown corresponding to fore and hindlimb of each crispant (#1-3). Deletions are indicated by dashes. PAM and microhomologous sequences are marked by red letters and underscores, respectively.

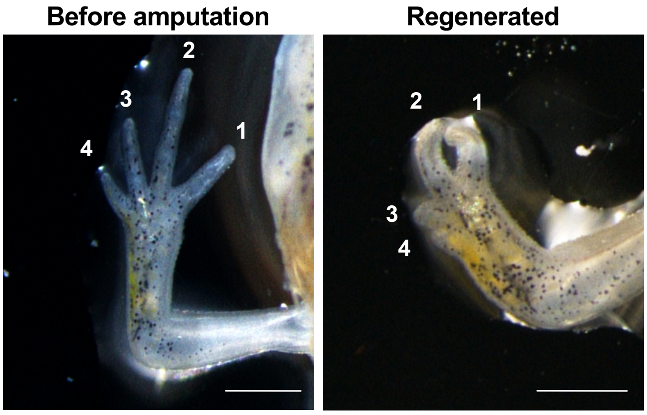


**Supplementary Fig. S5.** Forelimb regeneration of *Fgf10* crispant. Scale bar = 1 mm.

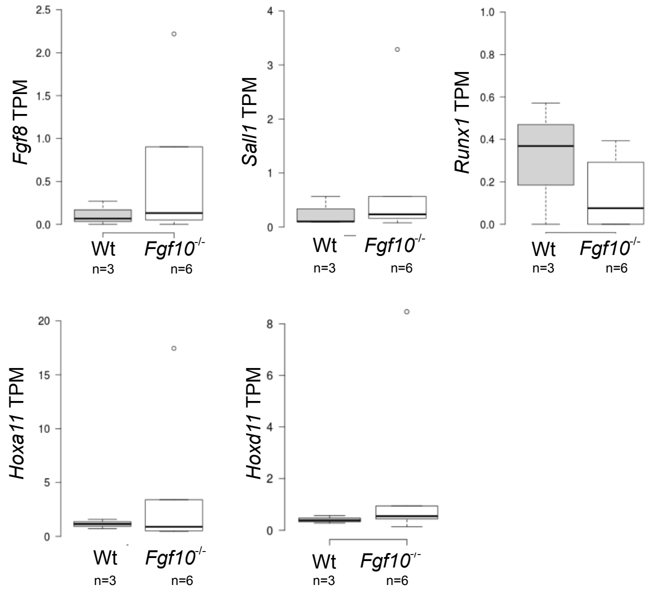


**Supplementaly Fig. S6.** Expression of limb development related genes in hindlimb blastema of wild type and *Fgf10* null mutant (n=3 Wt; 6 Fgf10 mutant; *p < 0.05).
